# Polymerase theta-helicase promotes end joining by stripping single-stranded DNA-binding proteins and bridging DNA ends

**DOI:** 10.1101/2021.06.03.446937

**Authors:** Jeffrey M. Schaub, Michael M. Soniat, Ilya J. Finkelstein

**Affiliations:** Department of Molecular Biosciences and Institute for Cellular and Molecular Biology, The University of Texas at Austin, Austin, Texas 78712, USA; Center for Systems and Synthetic Biology, The University of Texas at Austin, Austin, Texas 78712, USA

**Keywords:** SF2 helicase, Replication Protein A, RAD51, PARP-1, PARylation, single-molecule, DNA curtains

## Abstract

Homologous recombination-deficient cancers rely on DNA polymerase Theta (Polθ)-Mediated End Joining (TMEJ), an alternative double-strand break repair pathway. Polθ is the only vertebrate polymerase that encodes an N-terminal superfamily 2 (SF2) helicase domain, but the role of this helicase domain in TMEJ remains unclear. Using single-molecule imaging, we demonstrate that Polθ-helicase (Polθ-h) is a highly processive single-stranded DNA (ssDNA) motor protein that can efficiently strip Replication Protein A (RPA) from ssDNA. Polθ-h also has a limited capacity for disassembling RAD51 filaments but is not processive on doublestranded DNA. Polθ-h can bridge two non-complementary DNA strands *in trans.* PARylation of Polθ-h by PARP-1 resolves these DNA bridges. We conclude that Polθ-h removes RPA and RAD51 filaments and mediates bridging of DNA overhangs to aid in polymerization by the Polθ polymerase domain.

## Introduction

DNA double-strand breaks (DSBs) are highly toxic lesions that occur during cellular metabolism and in response to cancer therapies. Non-homologous end-joining (NHEJ)—the predominant DSB repair pathway in human cells—initiates when the ring-like Ku70/80 heterodimer binds the free DNA ends (1, 2). Subsequently, Ku recruits additional repair factors to the DSB, including DNA-PKcs to bridge the DNA ends and ligases to seal the break (3, 4). Homologous recombination (HR) is an error-free repair pathway that partially processes the free DNA ends to expose 3’-single-stranded DNA (ssDNA) overhangs (5). These overhangs are rapidly bound by replication protein A (RPA). Subsequently, RAD51 replaces RPA on the ssDNA to search for sequence homologies in a sister chromatid (6). RAD51-mediated strand invasion facilitates templated polymerization of a homologous DNA sequence (7). While NHEJ is active throughout the cell cycle, HR is restricted to the S and G2 phases of the cell cycle when a homologous template is available (8, 9).

Many cancer types accumulate mutations in NHEJ- or HR-dependent proteins and become reliant on theta-mediated end-joining (TMEJ), an error-prone DSB repair pathway (10). TMEJ is mediated by DNA Polymerase Theta (Polθ), PARP-1, and DNA Ligase III, as well as traditional DNA resection factors (11–13). Unlike HR, TMEJ requires short microhomologies (2-6 bp) and is highly mutagenetic, leading to increased chromosomal rearrangement and short insertions/deletions (14–16). During TMEJ, 3’-ssDNA overhangs are generated by MRE11-RAD50-NBS1 complex (MRN)/CtIP-mediated resection and are annealed at microhomologies (17). The resulting flaps are nucleolytically removed, and Polθ further extends the junctions to stabilize the microhomology (18). However, resected ssDNA is rapidly bound by RPA, which blocks the annealing of microhomologies (19). Furthermore, the recombinase RAD51 displaces RPA with the help of BRCA2 and other recombination mediators (20). Inactivation of TMEJ leads to an increase in HR, suggesting that these two repair pathways are antagonistic (21, 22)

Polθ is evolutionarily conserved across higher eukaryotes but is missing in fungi (23). Polθ encodes an N-terminal superfamily 2 (SF2) helicase/ATPase domain, a central disordered domain, and a C-terminal A-family polymerase domain (**Figure 1A**) (24, 25). The isolated Polθ-helicase (Polθ-h) domain is an ssDNA-dependent ATPase that can unwind short DNA duplexes and displace RPA from oligo-length DNA substrates *in vitro* (26, 27). In cells, Polθ-h mutants increase the prevalence of RAD51 foci after radiation exposure and shift the spectrum of end-joining products with microhomologies near the 3’ ends of DNA substrates (15, 21). This polymerase is frequently overexpressed in cancers deficient in traditional DSB repair mechanisms, and elevated Polθ expression led to poor patient prognosis (28–30). Inhibition of Polθ-h can kill HR-deficient tumor cells, suggesting a therapeutic route for targeting such malignancies (31). Polθ is an especially promising therapeutic target when combined with PARP-1 inhibitors in NHEJ/HR-deficient cancers (21, 22, 31–33). Together, these studies have established Polθ-h as a critical but enigmatic factor in TMEJ.

**Figure 1:**
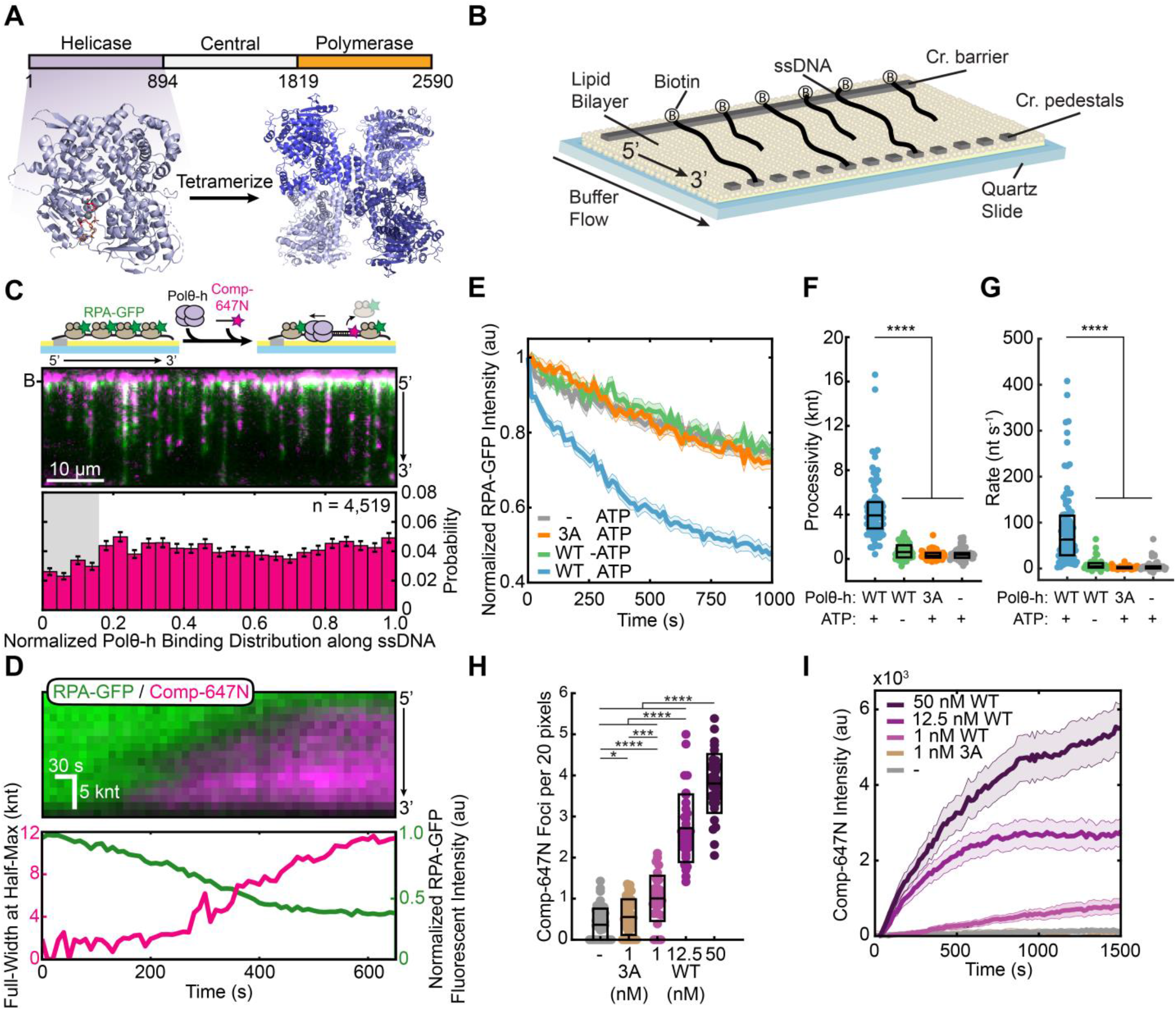
Polθ-h processively removes RPA from single-stranded DNA. **(A)** Polθ domain map (top) and a crystal structure of Polθ-h (PDB: 5A9J). **(B)** Schematic of ssDNA curtains assay. For double-tethered ssDNA curtains, buffer flow is stopped after DNA molecules are immobilized between the chromium barriers and pedestals. **(C)** Schematic (top) and microscope image (middle) of Polθ-h stripping RPA-GFP (green) from the ssDNA (top). The cleared regions are marked with a fluorescent complementary oligo (magenta). Histogram of where Polθ-h initiates RPA-GFP removal (bottom). **(D)** (top) Kymograph of processive RPA-GFP (green) removal, as measured by fluorescent complementary oligonucleotide (magenta). Bottom: Analysis of Polθ-h translocation (magenta) and RPA-GFP fluorescence intensity (green). **(E)** Normalized RPA-GFP fluorescent intensity for the indicated experimental conditions. Solid line (average), shading (±SEM). N > 46 for all conditions. **(F)** Polθ-h processivity and **(G)** velocity on RPA-coated ssDNA. Box displays median and IQR. N > 46 for all conditions. **(H)** Quantification of fluorescent oligonucleotide foci for each Polθ-h concentration on RPA-coated ssDNA. Box displays mean and S.D. **(I)** Fluorescent oligonucleotide intensity across each ssDNA for each Polθ-h concentration. Solid line (average), shading (±SEM).

Here, we use single-molecule and ensemble biochemical approaches to investigate Polθ-h. Polθ-h is a processive 3’ to 5’ ssDNA-binding motor and can readily displace RPA from ssDNA. Polθ-h can also partially disassemble RAD51 filaments, although this activity is much lower than its ability to remove RPA. Additionally, Polθ-h can bridge two DNA molecules that mimic resection intermediates *in trans* in a reaction that does not require ATP, suggesting that the homotetrameric assembly may tether two arms of a double-strand break during TMEJ. These DNA bridges were resistant to high salt, suggesting additional protein factors may be required for DNA dissociation. Therefore, we investigated the role of PARP-1 regulation of Polθ. PARP-1 rapidly binds to DNA damage sites and initiates the synthesis of poly ADP-ribose (PAR) chains on itself and client proteins that include Polθ. We show that PARP-1 PARylates Polθ-h *in vitro* and reduces the ssDNA binding affinity and promotes dissociation. We conclude that PARP-1 may regulate Polθ-h activity to promote DNA polymerization after the microhomology is established.

## Results

### Polθ-helicase strips RPA from single-stranded DNA

We purified and confirmed that the Polθ helicase domain (amino acids 1-894, referred to as Polθ-h) assembles into homotetramers via calibrated size exclusion chromatography consistent with previous studies (**Figure 1A, Figure S1**) (34). Next, we monitored single Polθ-h complexes using single-stranded DNA (ssDNA) curtains (**Figure 1B**) (35, 36). In this assay, ssDNA is generated by rolling-circle amplification of a repeating 28-nucleotide minicircle with low structural complexity (37, 38). The 5’ end of the primer includes biotin and the resulting ssDNA molecule is immobilized on the surface of a fluid lipid bilayer via biotin-streptavidin interactions. The ssDNA is then extended from the tether point via mild buffer flow.

We first assayed how Polθ-h counteracts RPA-coated ssDNA because RPA inhibits hybridization of heteroduplex oligos during TMEJ (19). We monitored the removal of fluorescent RPA-GFP because multiple fluorescent labeling strategies resulted in hypoactive Polθ-h (**Figure S2A**) (39). In this assay, ssDNA curtains are assembled with RPA-GFP. Next, unlabeled Polθ-h is added to the flowcell, and unbound protein is washed out. RPA clearance is observed following injection of fluorescent complementary oligonucleotide that can tile across the ssDNA substrate (Comp-647N) (**Figure 1C**). Injecting Polθ-h into the flowcell created a punctate pattern with reduced RPA-GFP signal and increased fluorescent oligonucleotide binding. RPA clearance and oligo binding required Polθ-h, suggesting that the helicase clears the ssDNA by removing RPA. Polθ-h cleared RPA along the entire ssDNA molecule, with a slight decrease at the 5’ end due to optical interference from the chromium barrier (**Figure 1C, Figure S2D**).

Next, we quantified RPA removal on double-tethered ssDNA curtains. RPA-ssDNA is tethered to downstream microfabricated chromium features and buffer flow is then stopped to observe protein dynamics on the double-tethered ssDNA in the absence of hydrodynamic forces. With Polθ-h and 1 mM ATP, all RPA-free regions expanded with a 3’ to 5’ polarity (N=91 Polθ-h molecules), consistent with other SF2-family helicases (27, 40) (**Figure 1D**). This RPA-cleared zone was rapidly hybridized by Comp-647N, indicating that Polθ-h created RPA-free ssDNA regions. RPA-GFP intensity decreased more rapidly in the presence of Polθ-h and 1 mM ATP than in negative control experiments without ATP or an ATPase-dead Polθ-h (E121A, D216A, and E217A; termed the 3A mutant) (21) (**Figure 1E**). We also observed small Comp-647N puncta when Polθ-h and/or ATP were omitted from the reaction. These foci were static throughout the experiment and likely represent locations where RPA-GFP is transiently displaced by excess Comp-647N (**Figure S2E**, **Table S1**). To estimate the processivity and rate of Polθ-h translocation, we fit the Comp-647N signal to a Gaussian function and calculated the full-width at half-max for each time point (**Figure S2B**). Polθ-h is a processive enzyme, clearing ~3.9 kilonucleotides (knt; IQR = 2.7-5.1 knt; N=91 Polθ-h molecules) of RPA-coated ssDNA with a median velocity of 63 nt s^-1^ (IQR = 28-117 nt s^-1^, N=91) (**Figure 1F,G**). Increasing Polθ-h concentration increased the number of Comp-647N foci per unit length and increased the total Comp-647N fluorescence intensity along the ssDNA substrate (**Figure 1H, I, Figure S2F, Table S2**). These results indicate that Polθ-h loads at multiple distinct positions along the ssDNA substrate. Taken together, we show that Polθ-h is a processive 3’ to 5’ ssDNA motor that uses ATP hydrolysis to strip RPA from ssDNA.

Polθ-h can unwind short duplex DNA molecules and DNA-RNA hybrids with limited processivity (27). Having observed processive ssDNA translocation, we next tested whether Polθ-h is also a processive helicase. We used a double-stranded DNA (dsDNA) substrate with 3’-ssDNA overhangs that mimic TMEJ resection intermediates. Helicase activity generates ssDNA that can be monitored via a growing RPA-GFP signal (**Figure S3A**) (41). However, the RPA-GFP intensity did not change when Polθ-h and ATP were added to the flowcell (**Figure S3B**). Although we cannot rule out limited helicase activity below our ~500 bp resolution, we conclude that Polθ-h is not a processive helicase on dsDNA (25, 27, 34).

### Polθ-h poorly disassembles RAD51 filaments

In addition to clearing RPA, Polθ has been proposed to antagonize HR by removing RAD51 filaments from ssDNA (21, 22). To test this hypothesis, we developed an assay to monitor Polθ-h-dependent RAD51 removal. RAD51 turnover on ssDNA is stimulated by its intrinsic ATPase activity but can be inhibited by adding Ca^2+^ to stabilize the pre-formed filament (42). However, Ca^2+^ also inhibits Polθ-h translocation on ssDNA (**Figure S4A**). Therefore, we used the ATPase-deficient RAD51(K133R) to stabilize RAD51 on ssDNA with ATP and Mg^2+^ in the reaction buffer (**Figure S4B, C**) (43). This mutation disrupts the Walker B ATPase motif, permitting ATP binding but not hydrolysis. We confirmed that RAD51(K133R) rapidly displaces RPA-GFP from ssDNA similarly to wild-type RAD51, albeit with a slightly longer nucleation phase (**Figure S4D**). As expected, RAD51(K133R) filaments are also more stable than WT RAD51 when challenged with RPA-GFP in the presence of Mg^2+^ and ATP (**Figure S4E**). In sum, RAD51(K133R) filaments assemble on ssDNA but remain stable in a buffer that also supports Polθ-h translocation.

We next tested whether Polθ-h can strip pre-formed RAD51(K133R) filaments from ssDNA. We first coated the ssDNA with RAD51(K133R) and then injected Polθ-h with a low concentration of RPA-GFP to visualize any ssDNA that is created during RAD51 removal (**Figure 2A**). In the presence of Polθ-h, the RPA-GFP puncta were ~2-fold brighter (N=53) than Polθ-h(3A) and when Polθ-h was omitted (N=41 and N=46, respectively) (**Figure 2B**). On RAD51(K133R)-coated ssDNA, the median Polθ-h processivity was 1.3 knt (IQR = 0.5-1.9 knt, N=53) and the velocity was 8 nt s^-1^ (IQR = 3-19 nt s^-1^, N=53) (**Figure 2C, D, Table S3**). Processivity was reduced 3-fold and the velocity was 8-fold slower with RAD51(K133R) as compared to RPA. In contrast to the RPA removal reaction, increasing Polθ-h concentration has only modest effects on the number of RPA-GFP foci per ssDNA (**Figure 2E, Figure S4F, Table S4**). The total RPA-GFP fluorescent intensity along the entire ssDNA substrate increased only ~2-fold above control experiments with inactive Polθ-h (**Figure 2F**). These results indicate that Polθ-h loads at gaps or junctions in the RAD51 filament to partially disassemble stabilized RAD51 filaments.

**Figure 2:**
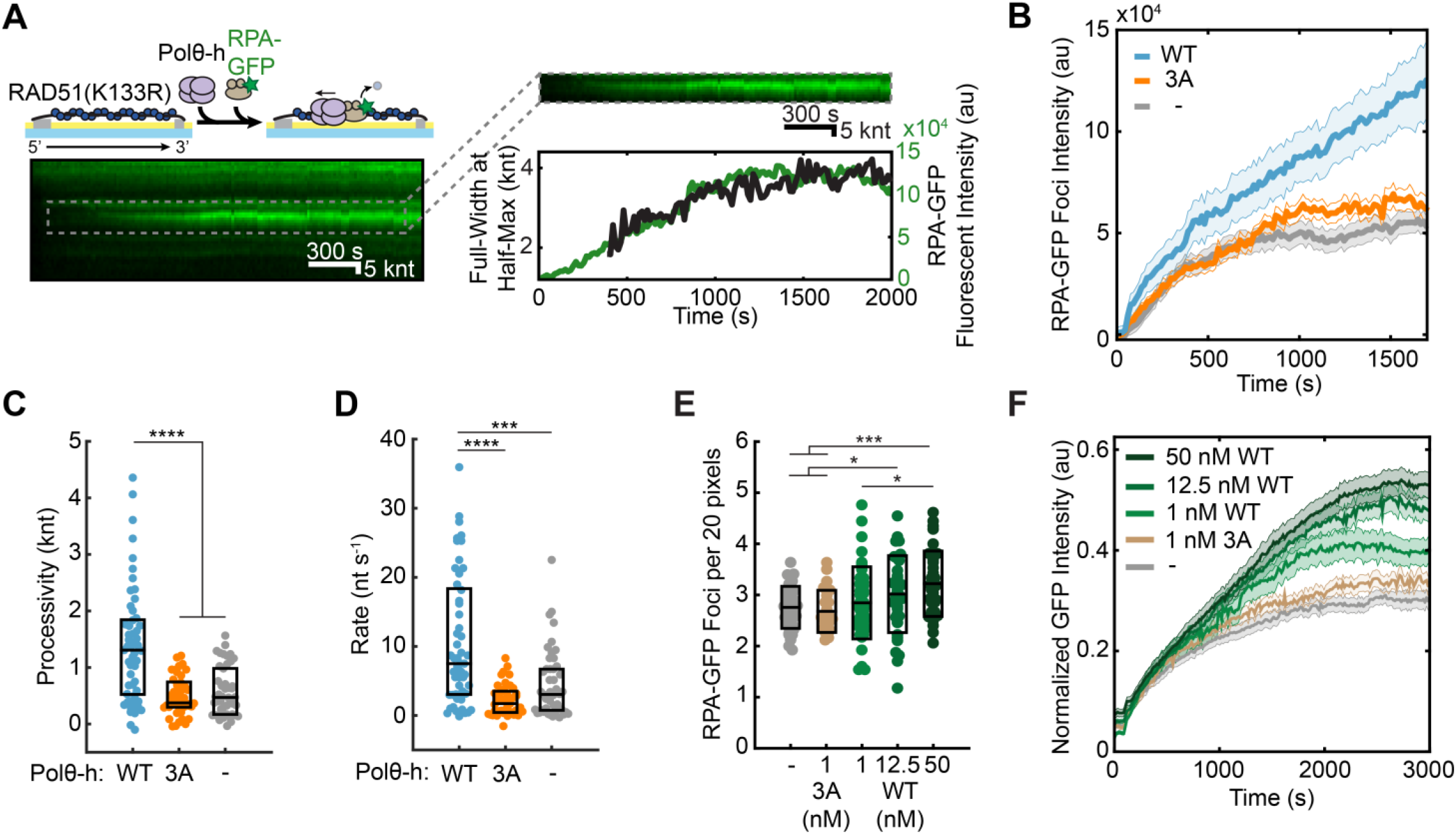
Polθ-h disassembles RAD51 filaments. **(A)** Cartoon, kymograph, and quantification of Polθ-h-mediated removal of RAD51(K133R). **(B)** Quantification of the RPA-GFP foci fluorescent intensity over time. Solid line (average), shading (±SEM). N > 36 for all conditions. **(C)** Polθ-h processivity and **(D)** velocity on RAD51(K133R)-coated ssDNA. Box displays median and IQR. N > 36 for all conditions. **(E)** Quantification of RPA-GFP foci for each Polθ-h concentration on RAD51(K133R)-coated ssDNA. Box displays mean and S.D. **(F)** Quantification of the total RPA-GFP intensity per DNA molecule at the indicated Polθ-h concentrations. Solid line (average), shading (±SEM). Time is normalized to when Polθ-h enters the flowcell (t=0). 2 nM RPA-GFP is immediately injected after, and flow is stopped at t=100s. Fluorescent intensity along ssDNA molecules is normalized to the initial RPA-GFP intensity prior to RAD51(K133R) displacement.

### PARP-1 reverses Polθ-h-mediated DNA bridges

TMEJ initiates after broken DNA ends are resected to reveal ssDNA overhangs (17). Polθ is proposed to bridge these overhangs despite their limited homology. The homotetrameric assembly of the helicase domain may underpin this multivalent DNA binding (34). To test whether Polθ-h can bridge thermodynamically unfavorable microhomologies, we first added Polθ-h to the ssDNA substrate and then flowed in a fluorescent non-complementary oligonucleotide (Noncomp-647N) (Figure 3A). Polθ-h efficiently captured this oligo, indicating that Polθ-h can bridge two ssDNA sequences regardless of homology. Notably, oligo capture required Polθ-h, whereas oligos did not associate with the ssDNA when Polθ-h was omitted (Figure S5A). We also tested whether Polθ-h can bridge DNA substrates that mimic DNA resection intermediates. We assembled 48 kbp-long dsDNAs with a 3’-T78 ssDNA overhang. Adding 5 nM Polθ-h resulted in bridging of adjacent molecules at their free DNA ends (Figure S5B). DNA bridging required Polθ-h but was ATPase independent; omitting ATP or using Polθ-h(3A) produced indistinguishable end-tethered DNAs (Figure S5C). These bridges persisted for the duration of the 15-minute imaging experiment. We additionally injected a ~60s pulse of 1M NaCl to attempt to dissociate Polθ-h from the ssDNA end. Surprisingly, end-tethering persisted through the high salt wash (Figure S5B).

**Figure 3:**
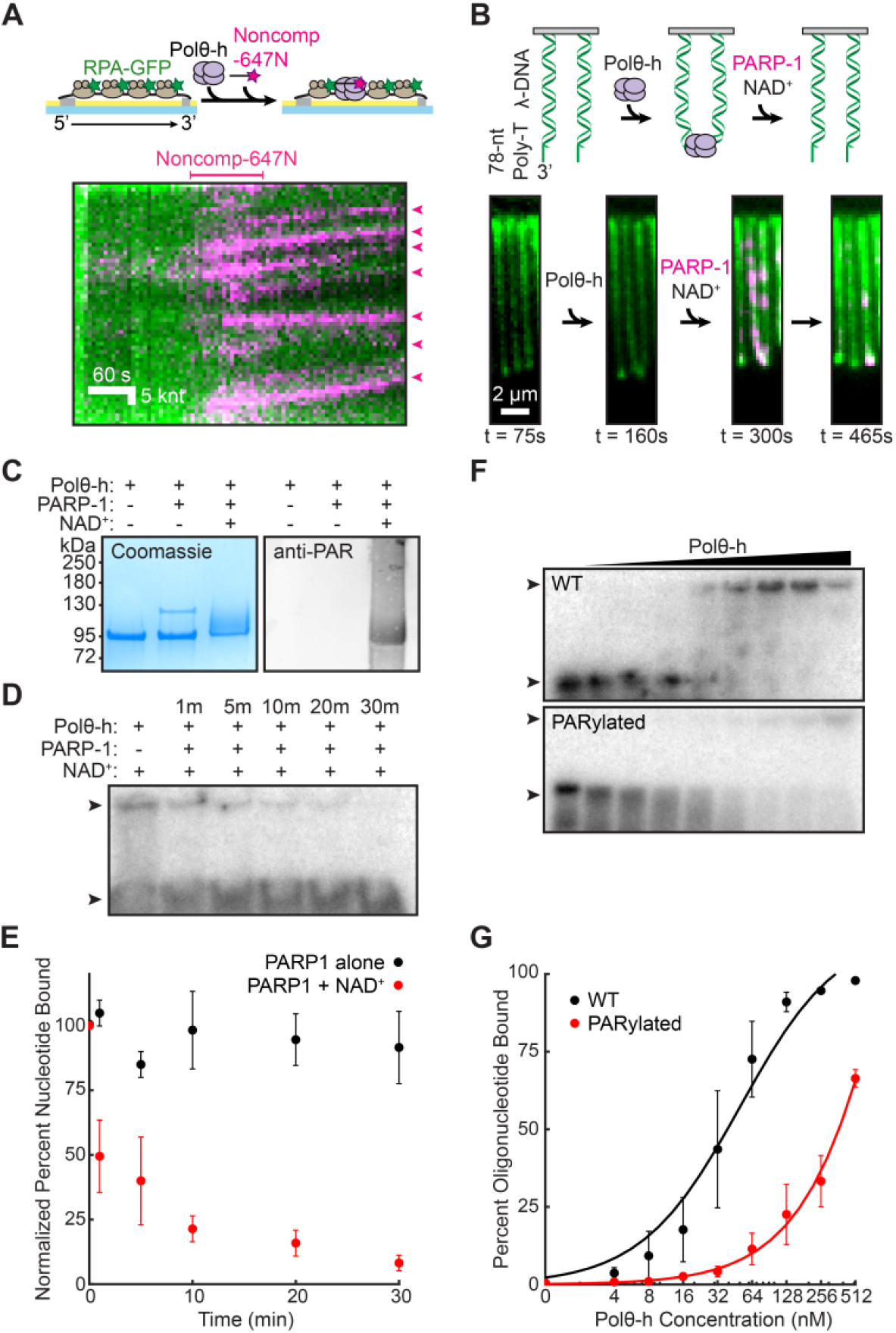
PARP-1 can resolve Polθ-h-mediated DNA bridges. **(A)** Cartoon (top) and kymographs (bottom) of Polθ-h-mediated tethering of two non-complementary ssDNA molecules. The long ssDNA is labeled with RPA-GFP (green) and the short non-complementary oligo is labeled with Atto647N (magenta). **(B)** Polθ-h tethers DNA molecules with 3’-ssDNA overhangs that mimic resected ends. Cartoon of DNA end bridging (top) and images of two tethered λ-phage DNA molecules. DNA is visualized with YOYO-1 (green). Addition of PARP-1 (magenta) and NAD^+^ dissociates these bridges. **(C)** Coomassie (left) and a western blot (right) indicating that Polθ-h is PARylated by PARP-1. **(D)** Electrophoretic mobility shift assay showing that ssDNA-bound Polθ-h dissociates from ssDNA after PARylation. Prebound Polθ-h and radiolabeled ssDNA oligonucleotide were incubated with PARP-1 and NAD^+^ for the indicated times. Arrows indicate unbound and bound oligonucleotide. **(E)** Quantification of (D). Binding normalized to condition without PARP-1. Average of three replicates. Error = SEM. **(F)** WT and PARylated Polθ-h EMSA on a radiolabeled ssDNA oligonucleotide. Polθ-h concentrations range from 0 to 512 nM. Arrows indicate unbound and bound oligonucleotide. **(G)** Quantification of (F). Fit to hyperbolic functions. Average of three replicates. Error = SEM.

We reasoned that Polθ-h-DNA bridges must be resolved for downstream TMEJ by another protein factor. PARP-1 is an attractive candidate for this activity for three reasons. First, PARP-1 is one of the earliest enzymes to arrive at broken DNA ends and plays a critical role in promoting TMEJ (13, 44, 45). Second, poly-ADP-ribosylation of client proteins by PARP-1 results in their release from DNA (46–49). Third, a proteomics screen identified the N-terminus of Polθ (i.e., the helicase domain) as a PARylation target (50). Consistent with our hypothesis, adding PARP-1 and NAD^+^ dissolved resected DNA bridges (Figure 3B). Omitting either PARP-1 or NAD^+^ was not sufficient to resolve these DNA bridges alone (Figure S5B). Purified PARP-1 can also PARylate Polθ-h *in vitro,* as indicated by a supershift of the Polθ-h SDS-PAGE band upon incubation with PARP-1 and NAD^+^ (Figure 3C, Figure S5D, E) An anti-PAR western blot confirmed that the upshifted Polθ-h band represents a PARylated product.

We further quantify whether PARylated Polθ-h has impaired ssDNA binding relative to the unmodified enzyme using electrophoretic mobility shift assays (EMSAs). Polθ-h was pre-incubated with a radiolabeled dT50 oligonucleotide prior to the addition of PARP-1 and NAD^+^. ssDNA-bound Polθ-h is rapidly released from ssDNA upon PARylation (Figure 3D, E). ssDNA remained bound by Polθ-h in the presence of only PARP-1 (Figure 3E, Figure S5F). We also changed the order of addition by pre-incubating Polθ-h with PARP-1 and NAD^+^ prior to incubating with a radiolabeled dT50 oligonucleotide (Figures 3F). Unmodified Polθ-h had a 39 ± 12 nM ssDNA binding affinity, which closely matches the 30 nM affinity measured via fluorescence anisotropy assays (34). In contrast, PARylated Polθ-h decreased ssDNA affinity at least ten-fold compared to Polθ-h alone (> 370 nM) (Figure 3G). Taken together, the single-molecule and ensemble experiments demonstrate that PARP-1 can PARylate Polθ-h and that PARylation reduces the ssDNA binding affinity of Polθ-h.

## Discussion

Figure 4 summarizes our model for how Polθ uses its unique helicase domain during TMEJ. Polθ encounters RPA-coated ssDNA that is generated during resection. Its helicase domain translocates in a 3’ to 5’ direction to processively remove RPA and other ssDNA-binding proteins from the ssDNA substrate. RPA prevents the hybridization of short microhomologies, so its removal is critical during TMEJ (19). Polθ-h removes RPA over thousands of nucleotides and can also partially disassemble RAD51 filaments. We used a RAD51 mutant that stabilizes ssDNA filaments in these studies, so these results are likely a lower estimate on Polθ-h’s ability to clear wild-type RAD51 filaments. We conclude that the helicase domain can load within RPA-coated segments or at RAD51-RPA filament junctions to rapidly remove RAD51 over the tens to hundreds of nucleotides that are required to synapse TMEJ junctions in cells (51–53). After clearing the ssDNA, Polθ-h bridges two DNA ends. This may be sufficient for the polymerase domain to extend the microhomologies. Following polymerization, these bridges can be resolved by PARP-1-dependent Polθ PARylation, which reduces the affinity of the enzyme for ssDNA. Removing Polθ may be required for ligases to re-seal the broken DNA breaks.

**Figure 4:**
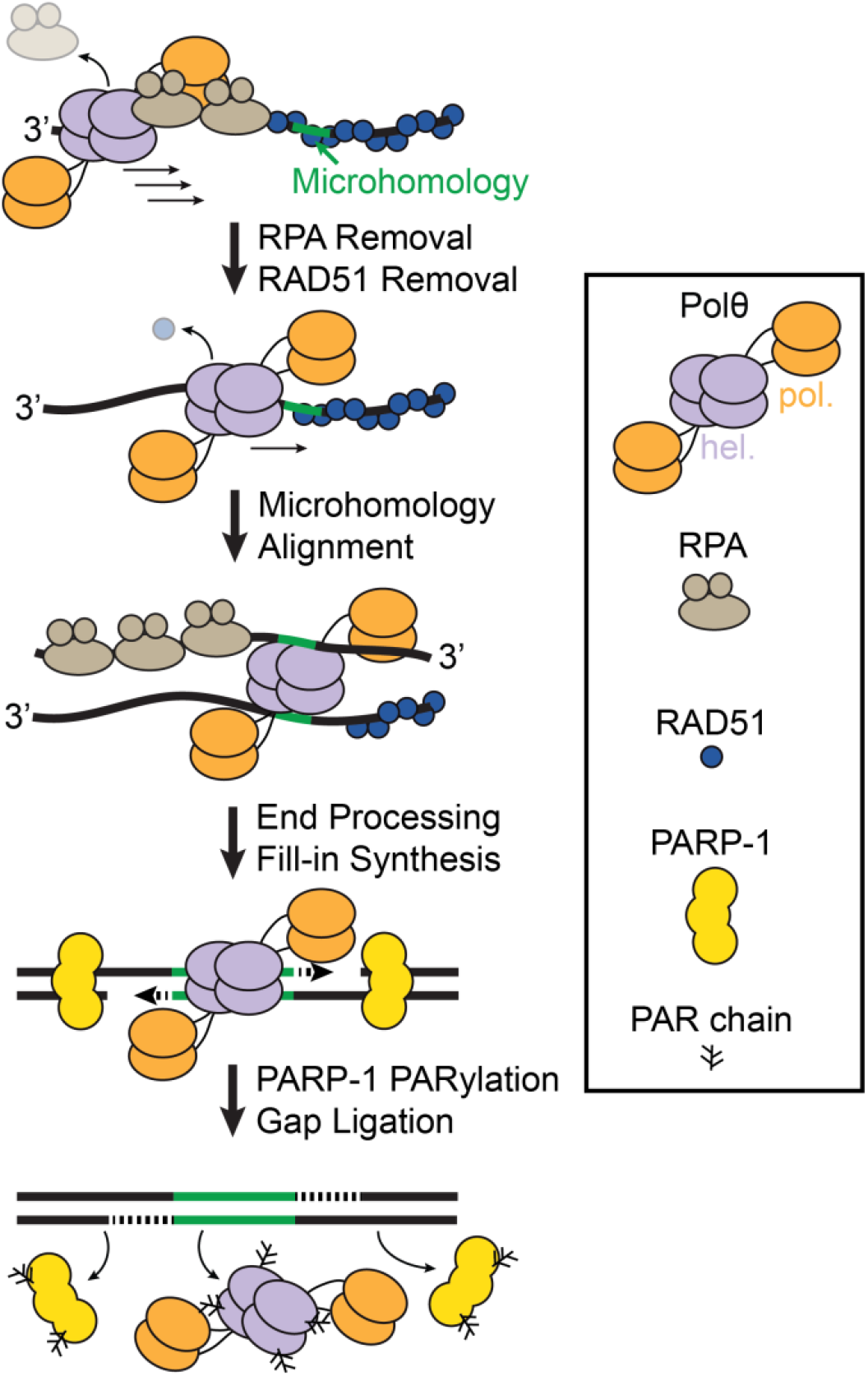
Model of Polθ activities in Theta-mediated end-joining. Polθ-h moves 3’ to 5’ on resected ssDNA to remove RPA and RAD51 and scans for microhomologies (green). 3’ ends are processed and resulting gaps are filled in by the Polθ polymerase domain. PARP-1 PARylates Polθ to remove it from DNA after gap filling.

By removing RPA, Polθ accomplishes the twin goals of exposing short microhomologies and suppressing the RPA to RAD51 exchange that precedes large RAD51 foci in cells. Indeed, Polθ ATPase mutants that disrupt the helicase domain shift the spectrum of TMEJ junctions to external microhomologies in cells. This result is presumably due to its inability to processively translocate and remove ssDNA-binding proteins to expose internal microhomologies (15, 51). We also observed that Polθ-h loads more efficiently on RPA-vs RAD51-coated ssDNA in a concentration-dependent manner. RPA rapidly exchanges and diffuse on ssDNA and we propose that this dynamic nature may promote Polθ-h’s loading on ssDNA compared to RAD51 filaments (54–56). Our observation that Polθ-h has limited RAD51 clearing activity is consistent with previous studies, including reports that RAD51 foci increase in cells that have helicase-dead Polθ (21, 22). The robust RPA removal activity that we observed biochemically suggests that RPA clearance is a major target for Polθ-h in cells.

The Polθ-h domain is postulated to be a reverse helicase, or annealase, that can thermodynamically hybridize short microhomologies (26). In this study, we show that Polθ-h can bridge two ssDNA sequences regardless of sequence homology. Based on this result, we suggest that the microhomology selection is mediated by the polymerase domain where Polθ-h initiates a 3’ to 5’ processive “microhomology scan” for the polymerase domain (15, 51). Surprisingly, these ssDNA bridges are highly resistant to NaCl, suggesting that additional protein factors are required for their disassembly.

PARP-1 is one of the first DNA damage sensing proteins to localize to DNA damage (57). Polθ recruitment to laser damage is reduced in cells with PARP inhibitors or PARP-1 depletion (22). Our data suggest that PARP-1 may further regulate the activity of Polθ beyond recruitment. PARP-1 binds with high affinity to DSB and ss/dsDNA junctions (58, 59). We propose that the PARylation activity on Polθ may aid in regulation and dissociation post-microhomology synthesis. Polθ binds to the resected 3’ ssDNA and processively translocases internally where PARP-1 then potentially PARylates Polθ. This may function in increasing the access to the polymerized DNA for ligation by the LIG3-XRCC1 complex (60). Additionally, PARylation may aid in the iterative microhomology selection and multiple rounds of DNA synthesis via regulation of Polθ DNA-binding (15, 61). We also do not rule out that PARylation by PARP-1 may inhibit Polθ DNA binding to favor more accurate forms of repair. Together, this work shows that Polθ plays multiple roles in mediating end-joining at DSBs in NHEJ/HR-deficient cancers and reiterates the importance of understanding the mechanistic functions of Polθ as a promising therapeutic target (12, 30–32).

## Materials and Methods

### Proteins and Nucleic Acids

Oligonucleotides were purchased from IDT or IBA (for fluorescent oligos). Polθ-h (amino acids 1-894) was cloned into a pET19 vector with an N-terminal 6xHis-TwinStrep-SUMO tag to generate pIF378. Polθ-h(3A) mutations E121A, D216A, and E217A were cloned with primers IF733 and IF734 using QuikChange Lightning Multi Site-Directed Mutagenesis Kit to generate plasmid pIF585 (Agilent #210516). RAD51(K133R) was mutagenized using inverse PCR with primers IF724 and IF725. RPA-GFP (plasmid pIF48), RAD51 (plasmid pIF224), RAD51(K133R) (plasmid pIF582) and PARP-1 (plasmid pIF662) were purified as previously described (41, 62, 63).

Polθ-h and Polθ-h(3A) were purified as described with some modifications (26). Plasmids were transformed into Rosetta(DE3) pLysS (Novagen) *E. coli* cells. Cell pellets were resuspended in Lysis Buffer (25 mM HEPES pH 8.0, 250 mM NaCl, 10 mM imidazole pH 8.0, 5 mM 2-mercaptoethanol, 10% glycerol and supplemented with Roche cOmplete protease inhibitor) and sonicated. The lysed pellet was centrifuged at 40,000 rcf for 45 minutes. Resulting clarified lysate was placed on a HisTrap column (GE Healthcare) and eluted on a gradient from 10 – 250 mM imidazole. Eluted material was digested with SUMO Protease for two hours at 4°C and diluted with 25 mM HEPES pH 8.0 to a final NaCl concentration of 100 mM. This was passed through a heparin column (GE Healthcare) and eluted with a gradient from 50 – 1000 mM NaCl. Pure Polθ-h eluted around 600 mM NaCl. Polθ-h-containing fractions were pooled, dialyzed in Dialysis Buffer (25 mM HEPES pH 8.0, 100 mM NaCl, 5 mM DTT, and 10% glycerol) for four hours at 4°C. Polθ-h was spin concentrated and flash-frozen in liquid nitrogen.

### Single-molecule Microscopy

Single-stranded DNA curtains were assembled in microfabricated flowcells according to published protocols (35, 36, 64, 65). All microscope experiments were conducted at 37°C. Images were collected on an inverted Nikon Ti-E microscope in a prism TIRF configuration running NIS Elements (AR 4.30.02). Flowcells were illuminated with a 488 and 637 nm lasers (Coherent OBIS) split with a 638 nm dichroic mirror (Chroma). Two-color images were recorded by twin electron-multiplying charge coupled device (EMCCD) cameras (Andor iXon DU897). Uncompressed TIFF stacks were exported from NIS Elements and further analyzed in FIJI (66). Data analysis was performed in MatLab R2019a (MathWorks).

### RPA Removal Assays

We first generated ssDNA in the flowcells as described previously (35, 65). Next, 0.4 nM RPA-GFP was added to Imaging Buffer (40 mM Tris-HCl pH 8.0, 2 mM MgCl2, 1 mM DTT, 0.2 mg mL^-1^ BSA, 50 mM NaCl, and 1 mM ATP) and injected at 0.4 mL min^-1^ to tether the ssDNA molecules at a chromium pedestal 13 μm away from the biotinylated anchors. Unbound RPA-GFP washed out with Imaging Buffer. Polθ-h was introduced at the indicated concentration at a flow rate of 0.4 mL min^-1^ and excess helicase flushed from the flowcell. To monitor Polθ-h activity, 2 nM complementary fluorescent oligo (Comp-647N) was added into the flowcell, and flow was stopped. Images with a 50 msec exposure were acquired every 15 seconds using a 14 mW 488 nm laser and 55 mW 637 nm laser (power measured at the front face of the prism). We fit the Atto647N fluorescent intensity along the DNA with a Gaussian function. The region of interest encompassed the ssDNA molecule and was three pixels wide to account for diffraction and ssDNA motion. The full-width at half-max (FWHM) of the Gaussian at each time point was used to measure the extent and rate of RPA-GFP clearance. For differing concentration injections of Polθ-h, foci were counted per unit length of the ssDNA molecule. For experiments that quantified total fluorescence intensity, we measured this intensity along the length of the entire ssDNA and normalized to a unit length to correct for heterogeneity in the ssDNA lengths.

### RAD51 Removal Assays

We first generated RPA-coated double-tethered ssDNA as described above. To assemble RAD51 filaments, 1 μM RAD51(K133R) was injected in Imaging Buffer supplemented with 1 mM CaCl2 and flow was stopped for 10 minutes. Flow was resumed at 40 μL min^-1^ to remove unbound RAD51. Polθ-h was introduced at the indicated concentration and a flow rate of 0.4 mL min^-1^. Because of RAD51’s strand capture activities, we could not use a fluorescent complementary oligo to monitor helicase translocation. Instead, we monitored RAD51(K133R) clearance by adding 2 nM RPA-GFP to the flowcell. At this concentration, RPA cannot readily replace RAD51(K133R) on the ssDNA. Images with a 50 msec exposure are acquired every 15 seconds using a 40 mW 488 nm laser. We fit the GFP fluorescent intensity to a Gaussian distribution. The FWHM of the Gaussian distribution at each time point measured the extent and rate of RAD51(K133R) clearance. Fluorescent molecules were quantified as described for the RPA clearance experiments described above.

### Polθ-h Helicase Assays

Polθ-h helicase activity was measured in flowcells containing double-stranded DNA (dsDNA), as used previously for RecQ-family helicases (41, 67). The DNA substrate was derived from bacteriophage λ. The *cosL* end was ligated with LAB07 and *cosR* with Lambda Poly-T that produces a 3’-T78 overhang (41). Polθ-h was injected into the flowcell in Imaging Buffer at 0.4 mL min^-1^. Unbound Polθ-h was washed out and the buffer was switched to Imaging Buffer containing 0.1 nM RPA-GFP at 0.4 mL min^-1^ to fluorescently label exposed ssDNA. The fluorescent intensity of RPA-GFP foci was calculated by averaging the area of a 3 x 3-pixel region of interest. We fluorescently stained DNA with YOYO-1 at the end of the experiment to confirm that RPA-GFP foci localized to DNA ends.

### DNA tethering assays

For single-stranded capture experiments, we first generated ssDNA as described above. 0.4 nM RPA-GFP is added to Imaging Buffer and flown through the flowcell at 0.4 mL min^-1^ to double-tether the ssDNA molecules. Unbound RPA-GFP was flushed out with Imaging Buffer and 1 nM Polθ-h was injected at 0.4 mL min^-1^. To monitor Polθ-h oligo capture, 2 nM noncomplementary fluorescent oligo (Noncomp-647N) was then added to the flowcell. Binding was monitored by acquiring 50 msec images every 15 seconds using 14 mW 488 nm laser and 55 mW 637 nm laser.

For double-stranded DNA end bridging experiments, we hybridized λ-phage DNA with LAB07 and Lambda Poly-T oligos by thermal melting and subsequent ligation with T4 DNA Ligase (NEB, M0202) as previously described (41). The DNA was fluorescently stained with YOYO-1 to visualize end-tethering.

For experiments involving PARP-1, 5 nM Polθ-h was injected in Imaging Buffer minus BSA. We omitted BSA here because Polθ-h PARylation was inhibited in the presence of BSA in our single-molecule experiments, possibly because BSA can act as a competitor for PARP-1 activity. PARP-1 was labeled with an anti-HA primary and goat antimouse QDot705 secondary antibodies (ICL RHGT-45A-Z and Thermo Q-11461MP) and injected into the flowcell. Next, we switched to Imaging Buffer supplemented with 50 μM NAD^+^ to initiate PARylation. End-tethering was monitored by acquiring 50 msec images every 5 seconds using 14 mW 488 nm laser.

### Ensemble PARylation

We performed Polθ-h PARylation reactions in automodification buffer (30 mM HEPES pH 8.0, 50 mM NaCl, 1.5 mM MgCl2, 1 mM DTT) with 1 μM Polθ-h, 500 nM PARP-1,4.5 mM NAD^+^, 500 nM annealed oligos (NJ061 and NJ062) at 30 °C (63). Western blots were imaged on an Odyssey imaging system (Licor) with anti-PAR primary and goat anti-mouse IR680 (Millipore Sigma AM80 and Abcam ab216776, respectively). A dT50 oligo was radioactively labeled with ^32^P by T4 PNK (NEB M0201). EMSAs were performed in Imaging Buffer at room temperature. Polθ-h ssDNA displacement EMSAs were performed in automodification buffer with 25 nM PARP-1.

## End Matter

### Author Contributions and Notes

J.M.S. and I.J.F. designed the experiments. J.M.S. performed the experiments and analyzed data. M.M.S. contributed to protein purification and data analysis. I.J.F. supervised the project. J.M.S., M.M.S., and I.J.F. wrote the manuscript.

## Acknowledgments

We thank Drs. Richard Pomerantz, Marc Wold, and Mauro Modesti for sharing over-expression vectors, and to members of the Finkelstein lab for carefully reading this manuscript.

## Data and materials availability

All data in the manuscript or the supplementary material is available upon request.

