## Supplemental Information for "Polymerase theta-helicase promotes end joining by stripping single-stranded DNA-binding proteins and bridging DNA ends"

### SUPPLEMENTAL MATERIAL

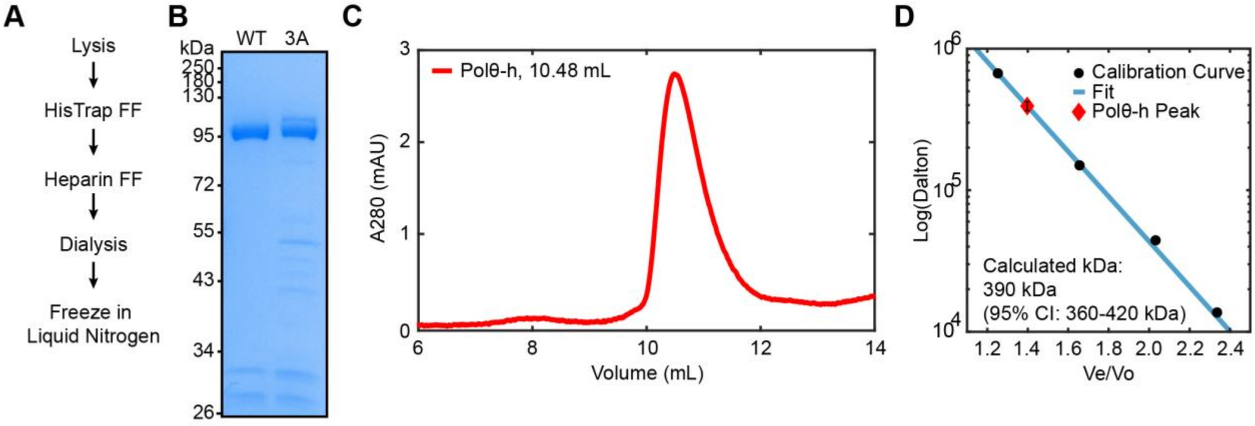


**Figure S1: Polθ-h purification and analysis. (A)** Schematic of the Polθ-h purification protocol. **(B)** SDS-PAGE gel of purified WT and 3A Polθ-h. The expected molecular weight is 99 kDa. **(C)** Superdex-200 chromatogram of recombinant homotetrameric Polθ-h. **(D)** Calculated molecular weight compared to a molecular weight standard (Sigma-Aldrich, 69385). Error bar within the diamond denotes a 95% confidence interval (CI).

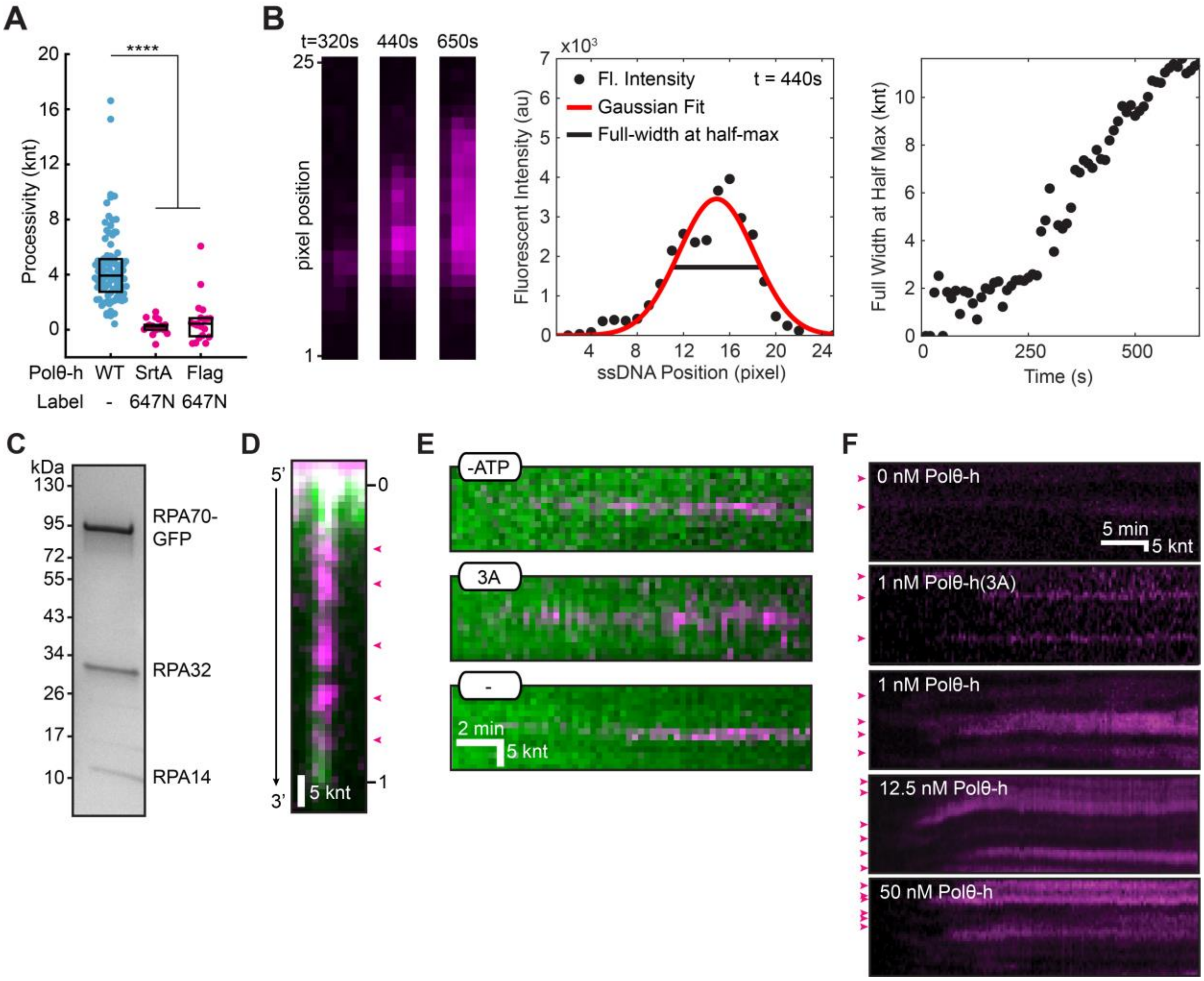


**Figure S2: Fluorescent analysis of Polθ-h variants and their translocation activities. (A)** We attempted to fluorescently label Polθ-h via sortase-mediated transpeptidation of the N-terminus or the linker region (C-terminus). N-terminal labeled Polθ-h was completely inactive. C-terminal Sortase (SrtA) or Flag epitope labeled Polθ-h constructs showed decreased processivity relative to WT Polθ-h on RPA coated ssDNA. Therefore, we focused on the WT Polθ-h for this study. Box displays median and IQR. **(B)** Summary of how Polθ-h translocation activity was analyzed via fluorescent proxies. Left: individual time points of fluorescent oligonucleotides hybridizing to the DNA. Middle: Gaussian fit of the resulting fluorescent signal. Right: A plot of the full width at half max of this fit as a function of time. **(C)** SDS-PAGE gel of recombinantly purified RPA-GFP. Expected molecular weights are 97, 29, and 14 kDa. **(D)** Analysis of Polθ-h RPA removal locations on single-tethered ssDNA curtains. Due to heterogeneity in their length, ssDNAs are normalized to unit length in a 5' to 3' direction. Magenta arrows show oligonucleotide foci that denote RPA removal activity. **(E)** Kymographs of Polθ-h-mediated RPA-GFP removal controls. ATP and hydrolysis activity are both required to see robust RPA removal. **(F)** Kymographs of concentration dependent Polθ-h oligonucleotide foci (magenta arrows) on RPA-GFP ssDNA curtains.

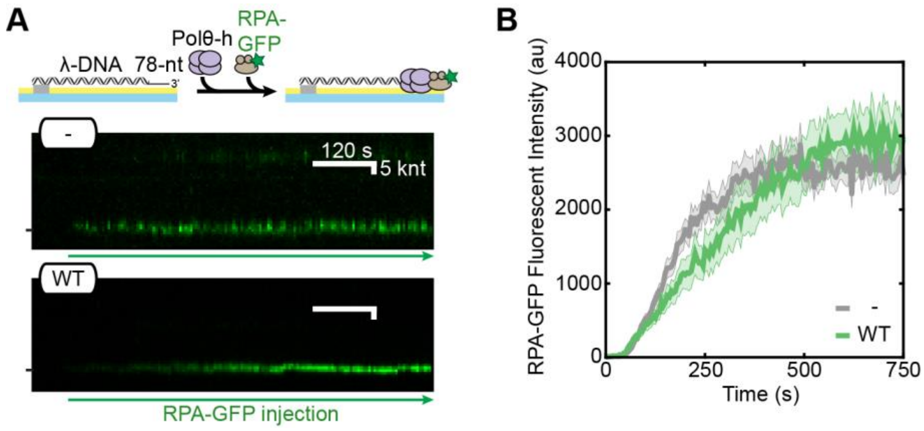


**Figure S3: Polθ-h is not a processive dsDNA helicase. (A)** Cartoon and kymographs of pre-resected helicase assay substrate in the presence and absence of Polθ-h WT. **(B)** RPA-GFP foci fluorescent intensity over time. WT (N=27) and without Polθ-h (-) (N=43).

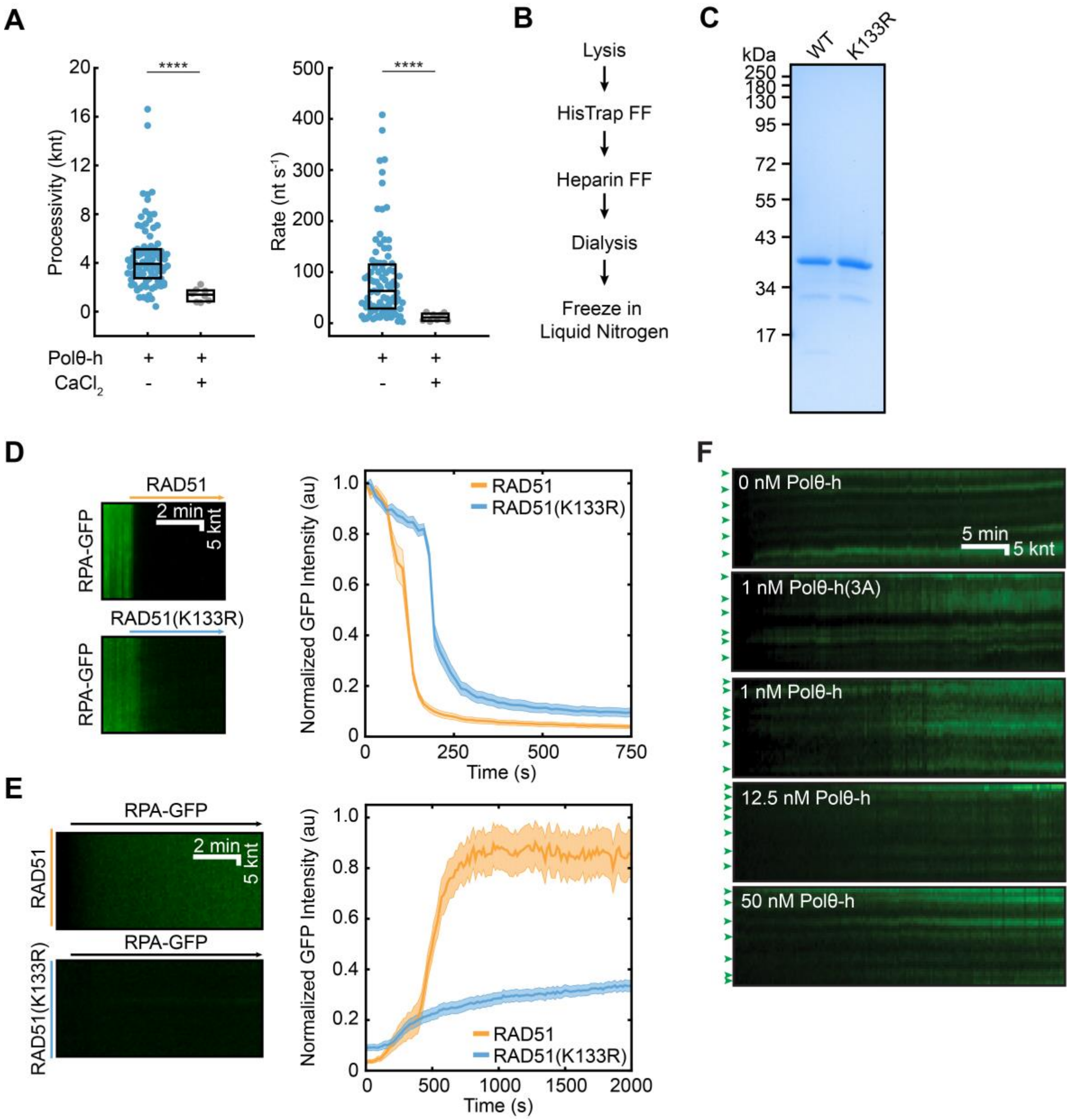


**Figure S4: Characterization of RAD51(K133R) and its exchange with RPA. (A)** Polθ-h is unable to remove RPA-GFP in the presence of $CaCl_2$. Boxes display the median and IQR. **(B)** Schematic of the RAD51 purification protocol. **(C)** SDS-PAGE gel of recombinant WT RAD51 and RAD51(K133R). Expected molecular weights are 37 kDa. **(D)** Kymographs of RAD51 dependent removal of RPA-GFP (left). Quantification of normalized RPA-GFP fluorescence over time (right). We analyzed 25 DNA molecules for both WT RAD51 and RAD51(K133R). **(E)** RAD51(K133R) is resistant to replacement by RPA as compared WT RAD51. Left: kymographs of RAD51 filament turnover as monitored by RPA-GFP. Right: quantification of the normalized RPA-GFP fluorescence (N=25 for both conditions). **(F)** Kymographs of RPA-GFP foci (green arrows) on RAD51(K133R) ssDNA curtains at the indicated Polθ-h concentrations.

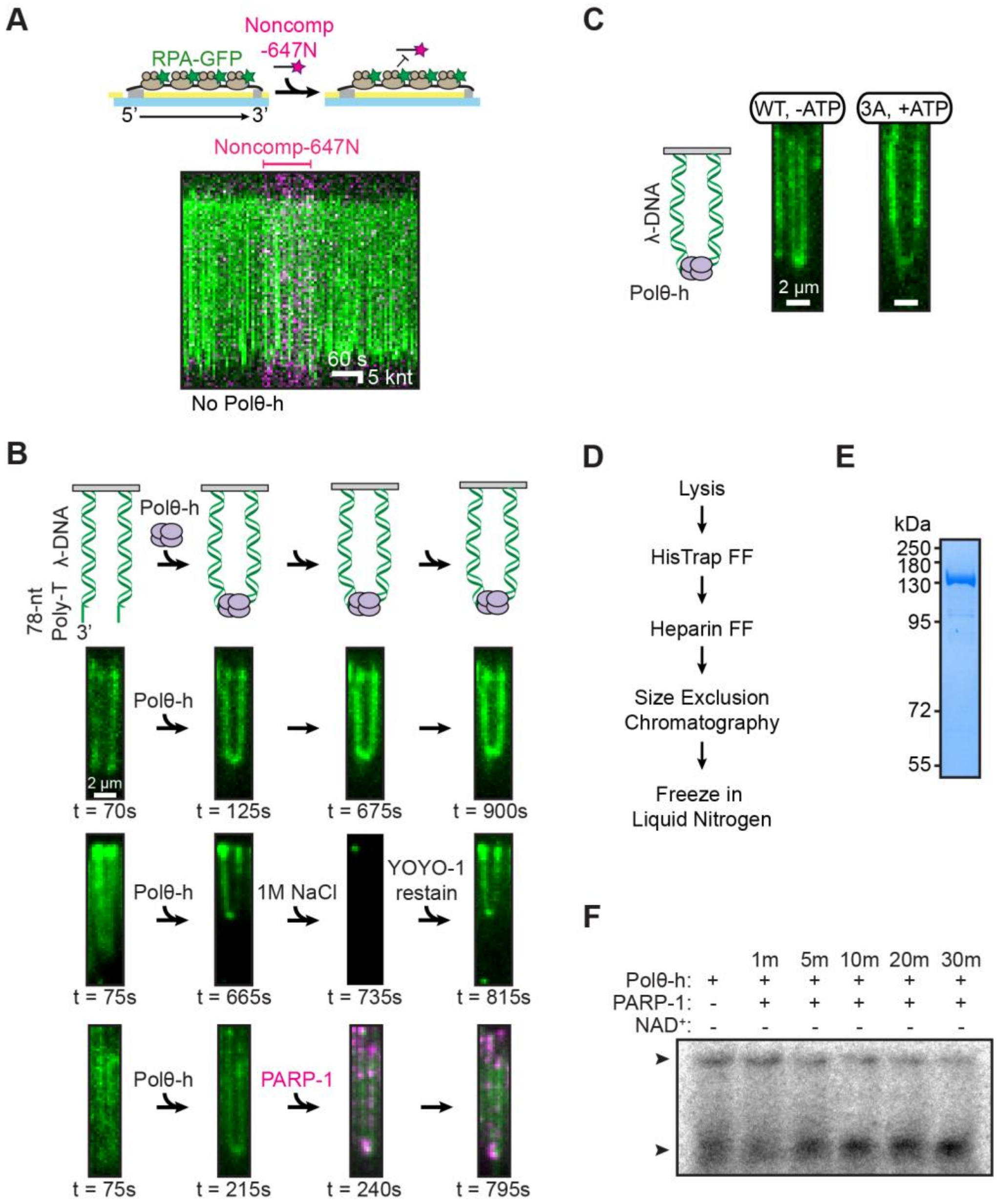


**Figure S5: DNA bridging requires Polθ-h and is ATP independent. (A)** Cartoon and kymograph of noncomplementary-647N (magenta) oligonucleotide injection in the absence of Polθ-h. ssDNA is bound with RPA-GFP (green). **(B)** Polθ-h tethering of two dsDNA ends persists for > 15 minutes and is resistant to 1M NaCl or the addition of PARP-1 (magenta) without $NAD^+$. The dsDNA substrate is visualized with the intercalating dye YOYO-1 (green). **(C)** Polθ-h tethers two ssDNA oligos annealed to a dsDNA molecule in the absence of ATP. The ATPase mutant, Polθ-h(3A), also tethered DNA. The dsDNA substrate is visualized with the intercalating dye YOYO-1 (green). **(D)** Schematic of the PARP-1 purification protocol. **(E)** SDS-PAGE gel of recombinantly purified PARP-1. N-terminal expression tags remain attached. Expected molecular weight is 129 kDa. **(F)** Prebound Polθ-h and radiolabeled ssDNA oligonucleotide were incubated with PARP-1 without $NAD^+$ over time.

**Table S1**: **Polθ-h velocity and processivity on RPA-ssDNA**

| Polθ-h | Nucleotide | Processivity, knt (IQR) | Relative processivity decrease | Rate, nt $s^{-1}$ (IQR) | Relative rate decrease | N (foci) |
|---|---|---|---|---|---|---|
| WT | ATP | 3.9 (2.7-5.1) | - | 63 (28-117) | - | 91 |
| WT | - | 0.6 (0.1-1.2) | 7x | 4 (2-12) | 16x | 57 |
| 3A | ATP | 0.3 (0.1-0.5) | 13x | 1 (0.5-4) | 63x | 46 |
| - | ATP | 0.4 (0.1-0.5) | 10x | 2 (0.2-5) | 32x | 55 |
| WT | ATP / $Ca^{2+}$ | 1.4 (0.8-1.8) | 3x | 11 (4-20) | 6x | 8 |
| SrtA-647N | ATP | 0.2 (0-0.4) | 20x | 5 (0-10) | 13x | 18 |
| Flag-647N | ATP | 0.4 (-0.5-0.8) | 10x | 10 (0-42) | 6x | 20 |

**Table S2**: **Quantification of Polθ-h foci on RPA-ssDNA**

| Polθ-h | Concentration, nM | Average foci per 20 pixels (S.D.) | N (ssDNA molecules) |
|---|---|---|---|
| WT | 50 | 3.8 (0.71) | 38 |
| WT | 12.5 | 2.7 (0.83) | 36 |
| WT | 1 | 1.0 (0.54) | 35 |
| 3A | 1 | 0.56 (0.45) | 41 |
| - | 0 | 0.37 (0.39) | 46 |

**Table S3: Polθ-h velocity and processivity on RAD51(K133R) filaments**

| Polθ-h | Nucleotide | Processivity, knt (IQR) | Relative processivity decrease | Rate, nt $s^{-1}$ (IQR) | Relative rate decrease | N (foci) |
|---|---|---|---|---|---|---|
| WT | ATP | 1.3 (0.5-1.9) | - | 8 (3-19) | - | 53 |
| 3A | ATP | 0.4 (0.3-0.6) | 3x | 2 (0-4) | 4x | 41 |
| - | ATP | 0.5 (0.2-1.0) | 3x | 3 (1-7) | 3x | 36 |

**Table S4: Quantification of Polθ-h foci on RAD51(K133R)-ssDNA**

| Polθ-h | Concentration (nM) | Average foci per 20 pixels (S.D.) | N (ssDNA molecules) |
|---|---|---|---|
| WT | 50 | 3.3 (0.63) | 38 |
| WT | 12.5 | 3.0 (0.75) | 40 |
| WT | 1 | 2.9 (0.70) | 43 |
| 3A | 1 | 2.7 (0.43) | 27 |
| - | 0 | 2.8 (0.41) | 44 |

**Table S5**: **Oligonucleotides used in this study.**

| Oligonucleotide | Sequence (5' to 3') |
|---|---|
| Template | /Phos/AG GAG AAA AAG AAA AAA AGA AAA GAA GG |
| Primer | /Biosg/TC TCC TCC TTC T |
| Comp-647N | /AT647/AGG AGA AAA AGA AAA AAA GAA AAG AAG G |
| Noncomp-647N | /AT647/TCC TCT TTT TCT TTT TTT CTT TTC TTC C |
| Poly-$T_{50}$ | $T_{50}$ |
| NJ061 | GGG TTG CGG CCG CTT GGG |
| NJ062 | CCC AAG CGG CCG CAA CCC |
| LAB07 | /Phos/AGG TCG CCG CCC/BioTEG/ |
| Lambda Poly-T | /Phos/GGG CGG CGA CCT $T_{78}$ |
| IF724 | GGA GAA TTC CGA ACT GGG AGG ACC CAG ATC TGT CAT ACG C |
| IF725 | GCG TAT GAC AGA TCT GGG TCC TCC CAG TTC GGA ATT CTC C |
| IF733 | CTC CTA CAA GTG CTG GGG CGA CTC TTG TGG CAG |
| IF734 | GGA ATG GTG GTT GTG GCT GCA TTA CAT ATG CTG GGA GAC TC |
